# Polymyxin heteroresistance in *Klebsiella oxytoca*

**DOI:** 10.1101/2025.08.01.668101

**Authors:** Eunice A Ayerakwa, Edward J.A. Douglas, Gerald Larrouy-Maumus, Andrew M. Edwards, Abiola Isawumi

## Abstract

Antibiotic heteroresistance presents a growing public health concern since the phenotype is associated with treatment failure and is hard to detect using conventional diagnostic testing. In this study, we characterized polymyxin B heteroresistance in a collection of clinical and environmental isolates of *Klebsiella oxytoca*, an opportunistic pathogen associated with hospital-acquired infections. All six of the isolates tested exhibited heteroresistance, indicated by a 16-fold difference between the highest non-inhibitory concentration (2 µg/mL) and the minimum inhibitory concentration (32-64 µg/mL) observed in population analysis profile (PAP) assays. Heteroresistance was found to be due to the presence of a stable sub-population of resistant bacteria and was unaffected by growth phase or the presence of host antimicrobial factors present in human serum. MALDI-TOF analysis revealed L-Ara4N modifications of the lipid A of resistant sub-populations. This pilot study identifies polymyxin heteroresistance in *K. oxytoca* may complicate treatment of infections caused by this organism.

## Introduction

Carbapenem- and third-generation cephalosporin-resistant pathogens including *Klebsiella pneumoniae, Escherichia coli*, and *Enterobacter* spp. have been classified as a critical priority group by the WHO (WHO BPPL, 2024). In addition, other species of the family Enterobacteriaceae are emerging as public health threats due to their ability to cause severe infections and resist multiple antibiotics. One such pathogen is *K. oxytoca*, the second most common cause of *Klebsiella* infection after *K. pneumoniae* (Singh et al., 2016), with some strains producing extended-spectrum beta-lactamase (ESBL), carbapenemase, imipenemase and/or oxacillinase enzymes (Yang et al., 2022; Shaikh et al., 2015; Ikhimiukor et al., 2023)). Additional resistance mechanisms include efflux pumps that confer intrinsic low level resistance to β-lactams and quinolones respectively (Yang et al., 2022). Moreover, several *K. oxytoca* isolates have acquired resistance to colistin, aminoglycosides, fosfomycin, sulphonamides, trimethoprim and macrolides (Yang et al., 2022). In keeping with similar pathogens, multiple drug resistance is often mediated by the presence of plasmids that encode diverse AMR determinants (Wang et al., 2017).

In humans, *K. oxytoca* is routinely found in the respiratory and gastrointestinal tract as part of the normal flora (Yang et al., 2022). However, the pathogen can establish mild to life-threatening infections including pneumonia, UTIs, bacteraemia, meningitis and antibiotic-associated haemorrhagic colitis in immune compromised individuals (Farooq et al., 2015; Youn et al., 2018). Additionally, *K. oxytoca* has been associated with skin and soft tissue infections such as surgical wounds, and necrotizing fasciitis in patients with malignancies and abscesses (Yang et al., 2022). In clinical settings, drug-resistant *K. oxytoca* has been responsible for infection outbreaks, most likely due to its ability to survive on fomites and hospital infrastructure such as water reservoirs, ventilators, sinks and disinfectants (Schmithausen et al., 2019). The emergence and dissemination of these resistant isolates has restricted treatment options, especially in resource-limited settings, necessitating the use of second- or third-line options.

Polymyxins are cationic lipopeptide antibiotics used as a last resort for treating infections caused by multi-drug resistant Gram-negative bacteria (Ledger et al., 2022). These antibiotics target lipopolysaccharide, disrupting both the outer and inner-membranes leading to bacterial death (Sabnis et al., 2021; Mohapatra et al., 2021). While polymyxins remain a treatment of last resort for highly drug resistant Gram-negative pathogens, there is growing concern about the frequency of polymyxin resistance (Li et al., 2019).

Resistance can arise spontaneously or via the acquisition of a mobile colistin resistance (*mcr*) gene. In both cases, resistance is conferred by modification of the lipid A component of LPS (da Silva et al., 2022). There is also a growing awareness of polymyxin heteroresistance (Ledger et al., 2022; Isawumi et al., 2020), which describes a population-wide variation in antibiotic response among bacterial subpopulations. Generally, two forms of antibiotic heteroresistance have been described; inducible and classical heteroresistance. Inducible heteroresistance may be triggered by antibiotic exposure such that transient phenotypic resistance occurs (Dewachter et al., 2019). Conversely, classical heteroresistance describes a stable subpopulation of highly resistant bacteria, usually due to mutations or gene amplifications (Nicoloff et al., 2019).

Antibiotic heteroresistance has been implicated in persistent infection and antibiotic therapeutic failure (Band & Weiss, 2019). This challenge is exacerbated by the difficulties associated with detecting heteroresistance using standard antibiotic susceptibility testing (Hassall et al., 2024). The current gold standard for heteroresistance detection is population analysis profiling (PAP) which involves quantifying the ability of a bacterial population to grow across a gradient of antibiotic concentrations (El-Halfawy & Valvano, 2015). The population is considered heteroresistant if the inhibitory concentration for the resistant sub-population is at least eight-fold higher than the minimum inhibitory concentration of the bulk population (Wan et al., 2023). However, this approach is time and resource demanding and is not typically used in diagnostic laboratories.

Given the growing importance of polymyxin antibiotics for treating infections in low income settings, and the challenges associated with diagnostics, the emergence of polymyxin heteroresistance in clinical isolates of *K. oxytoca* (Isawumi et al., 2020) signifies a major public health threat. Therefore, the aim of this study was to identify and characterise polymyxin heteroresistance in clinical and environmental *K. oxytoca* isolates from Ghana.

## Materials and Methods

### *K. oxytoca* isolates and growth conditions

Six *K. oxytoca* isolates a Ghanaian hospital ICU were sourced from the ABiola ISAwumi (ABISA) strain collection at West African Centre for Cell Biology of Infectious Pathogens, University of Ghana with ethical approval from the Ghana Health Service (GHS-ERC01/02/17). This included three clinical isolates (*Kleb401, Blood4a, Nasal2a)* from patients with bacteraemia, and three environmental isolates (*ACN, Kleb405, CTO4*) from ICU fomites (door handles and faucets). The isolates were selected based on their high levels of resistance to conventional and last resort antibiotics, including carbapenems. All isolates were grown in Mueller Hinton broth (MHB) at 37 °C with shaking at 180 RPM.

### Antimicrobial susceptibility profiling

The susceptibility profile of the strains to commonly used and last resort antibiotics in healthcare settings (ceftazidime, levofloxacin, chloramphenicol, gentamicin, imipenem, meropenem, colistin, trimethoprim, and polymyxin B) was determined using the broth microdilution method (Yang et al., 2022). Briefly, the antibiotics were serially diluted 2-fold across a 96-well plate with concentrations ranging from 64 µg/mL to 0.125 µg/mL. 100 µL of overnight cultures standardized to provide 5 × 10^5^ CFU/well was added to each well containing 100 µL of the antibiotics and incubated at 37 °C for 24 h. The minimum inhibitory concentration (MIC) of the antibiotics were determined by absorbance measurement at 600 nm using a microplate reader (TECAN, Infinite 200 Pro). The MIC was defined as the concentration of antibiotic that inhibits bacterial growth by >90% (no visible growth by eye).

### Population analysis profiling (PAP)

PAP assays were performed using log or stationary phase cultures following a previously described protocol (Wan et al., 2023). Briefly, a single colony of each isolate was inoculated into 3 mL of MHB and incubated overnight at 37 °C with shaking (180 RPM). For some assays, human serum was included at 50% culture volume to understand if this host mimicking condition influenced the heteroresistance profile. Bacteria were then used for PAP analysis (stationary phase) or, for log phase cultures, bacteria were sub-cultured and grown for 3 h with shaking. Seven serial dilutions (10^−1^ 10^−7^) were prepared from the cultures using PBS as the diluent. Ten microliters of each dilution was then spread on MH agar plates containing polymyxin B at a range of different concentrations (128, 64, 32, 16, 8, 4, 2 or 1 µg/mL). The plates were incubated at 37 °C for 24 h. The colony-forming unit on each MH agar plate was then used to determine CFU/ml by taking into account the dilution factor. Heteroresistance was identified if the highest inhibitory concentration at which there was no growth was ≥ 8-fold higher than the highest non-inhibitory concentration (concentration at which ≥ 80% of the bacterial population grew compared to the inoculum).

### Antibiotic killing assays

A single colony of each isolate was cultured in 3 mL of MHB overnight at 37 °C with shaking (180 RPM). MHB (3 ml) containing 4 µg/mL of polymyxin B was then inoculated with bacteria to 5 × 10^5^ CFU mL^−1^ and incubated at 37 °C with shaking (180 RPM). Aliquots (50 µL) of the cultures were recovered at specific time points (0, 2, 4, 6 and 24 h), serially diluted (10^1^ - 10^7^) in PBS and then plated on MH agar plate at different time points. The plates were incubated at 37 ° C for 24 h and CFU counts determined as described above.

### Serum susceptibility assays

Isolates (5 × 10^5^ CFU/mL) were incubated at 37 °C for 24 h in serial 2-fold dilutions of human serum (from 50% to 0.78% diluted in MHB) to determine their serum resistance profile. Bacterial growth was determined by measuring absorbance at 600 nm with a microplate reader (TECAN, Infinite 200 Pro), with values corrected against the relevant concentration of serum/MHB without bacteria.

### Time-kill assay in human serum

Serum time-kill assays were performed by inoculating 1 mL of human serum with bacteria to 5 × 10^8^ CFU mL^−1^ and incubated at 37 °C with shaking. Aliquots of the cultures were taken at specific time points (0, 30, 60, 90 and 120 min), serially diluted (10^1^ - 10^7^) in PBS and plated on MH agar. The plates were incubated at 37 °C for 24 h and the number of colony forming units was determined as described above.

### Lipid A analysis

Lipid A analysis was performed using the optimized MALDIxin test as described by Dortet et al., (2020). Briefly, 50 µL of bacterial culture was centrifuged, and pellets were washed three times in ultra-pure water. The pellets were subjected to mild-acid hydrolysis by resuspension in 2 % acetic acid and heating for 30 min at 100 °C. The acid treated cells were recovered by centrifugation and pellets washed twice and suspended in 50 µL of ultra-pure water. The bacterial suspension (0.5 µL) was loaded onto the target and mixed with 0.5 µL of a super-2,5-dihydroxybenzoic acid matrix. The mixture was air-dried, followed by MALDI-TOF analysis on a 4800 Proteomics analyzer (Applied Biosystems, Foster City, CA, USA).

## Results

### Antimicrobial susceptibility profile of K. oxytoca isolates

All six isolates (*CT04, Nasal2a, kleb405, Blood4a, ACN kleb401*) were susceptible to ceftazidime, gentamicin and meropenem with MICs ranging from 0.125 µg/mL to 2 µg/mL **(Supplementary Table S1)**. The isolates were resistant to levofloxacin (≥ 32 µg/mL), chloramphenicol (>64 µg/mL), imipenem (≥ 16 µg/mL), and trimethoprim (> 64 µg/mL). The MICs of colistin and polymyxin B for all six isolates ranged from 0.25 µg/mL to 64 µg/mL, with a skipped well phenotype suggestive of heteroresistance (**Figure 1**).

**Fig. 1:**
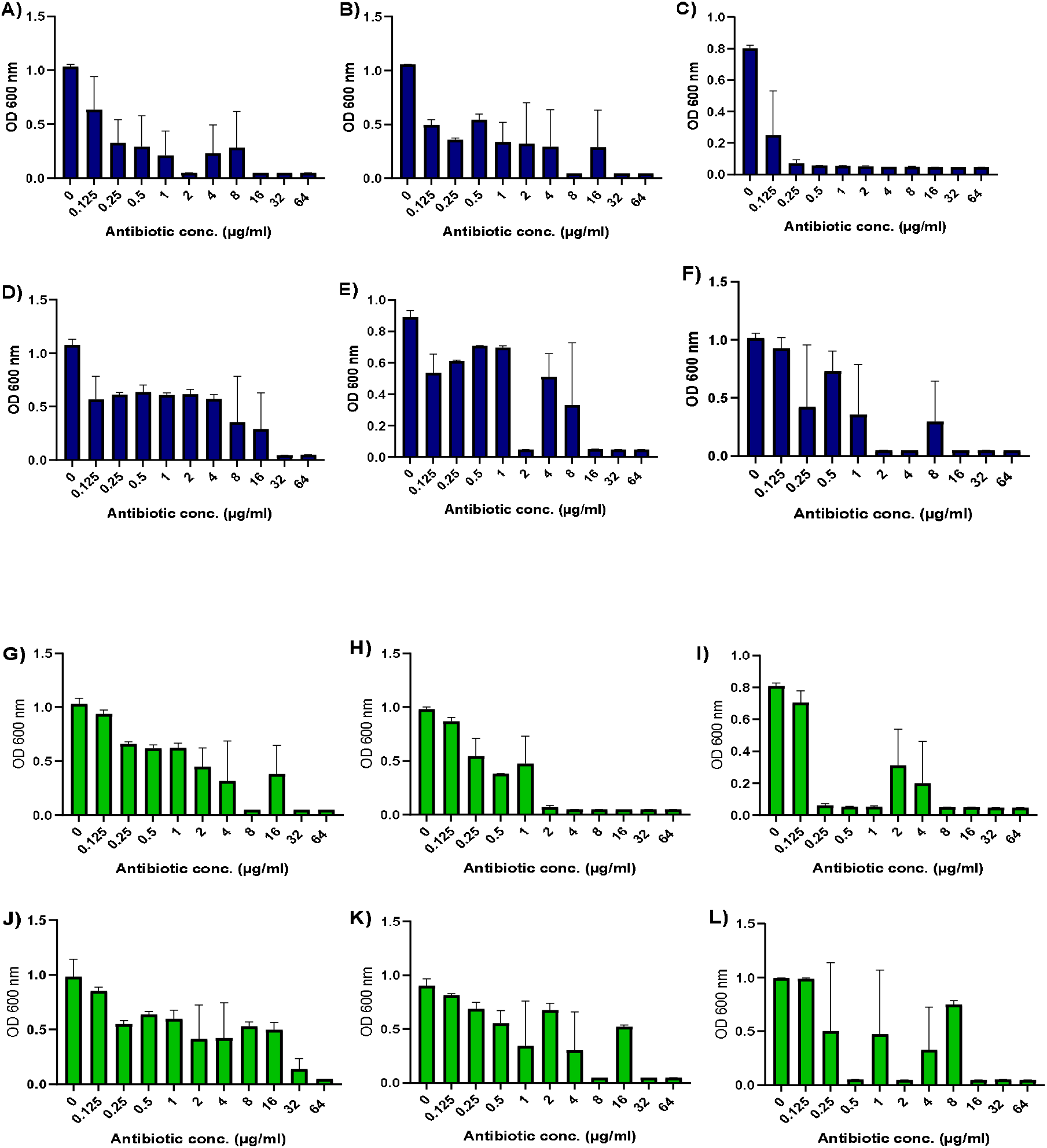
Growth (OD_600_) of strains in the presence of various concentrations of **Polymyxin B** (**A**. CT04, **B**. Nasal2a, **C**. Kleb405, **D**. Blood4a, **E**. ACN and **F**. Kleb401), and **Colistin** (**G**. CT04, **H**. Nasal2a **I**. Kleb405, **J**. Blood4a, **K**. ACN and **L**. Kleb401). The data represent three independent broth microdilution assays in MHB.

### Phenotypic characterization of polymyxin B heteroresistance in K. oxytoca

Polymyxin heteroresistance was confirmed in all six isolates using the PAP assay **(Figure 2)**. In each case, it was demonstrated that the MIC of the resistant sub-population (≥ 64 µg/mL) was 8-16 fold higher than their maximum non-inhibitory concentrations of the bulk population (2 µg/mL). However, the size of the sub-population varied between strains from ∼10^−6^ to 10^−7^. To determine the stability of the heteroresistance phenotype, a single colony of each resistant subpopulation (*CT04*^*R*^, *Nasal2a*^*R*^ *Kleb405*^*R*^, Blood4a^R^ *ACN*^*R*^, and *Kleb401*^*R*^) was picked from agar plates containing polymyxin B (32 µg/mL) and re-assayed using three independent PAP experiments. All six of the isolates showed stable homogenous resistance, with MICs >128 µg/mL, the maximum concentration used in the assay (**figure 2**).

**Fig. 2:**
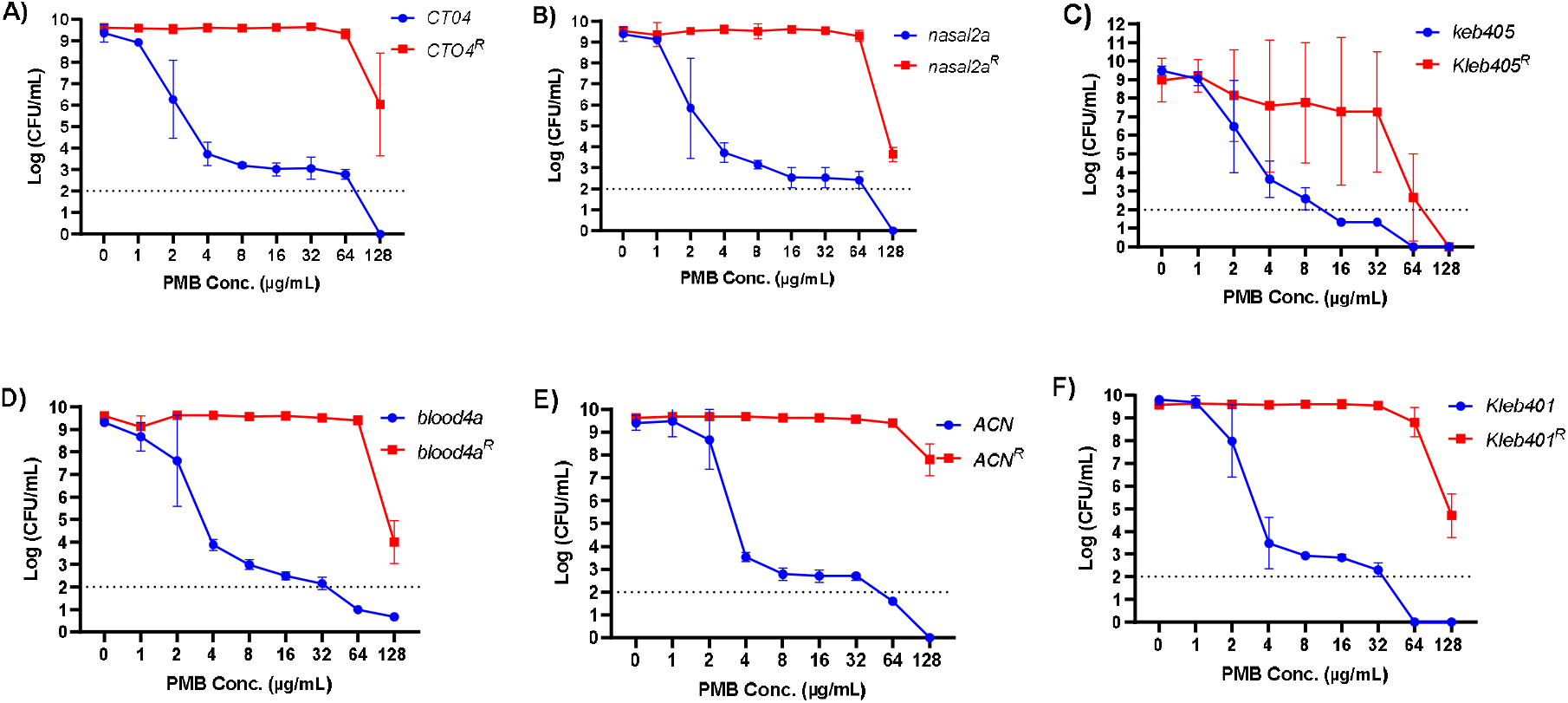
Polymyxin heteroresistance in *K. oxytoca* isolates. Graphs show the data from PAP assays for the six test isolates *A) CTO4, B) Nasal2a, C) Kleb405, D) Blood4a, E) ACN and F) Kleb401*. For each isolate, a single colony of resistant subpopulation (*CT04*^R^, *Nasal2a*^R^ *Kleb405*^R^, Blood4a^R^ *ACN*^R^, and *Kleb401*^R^) was re-assayed using PAP experiments to determine the stability of the phenotype. Three independent biological replicates (N=3) were performed and the log of the mean±SEM CFUs were plotted. The detection limit for viable colonies was 100 CFU/mL corresponding to log (2) on the y-axis.

**Fig. 3:**
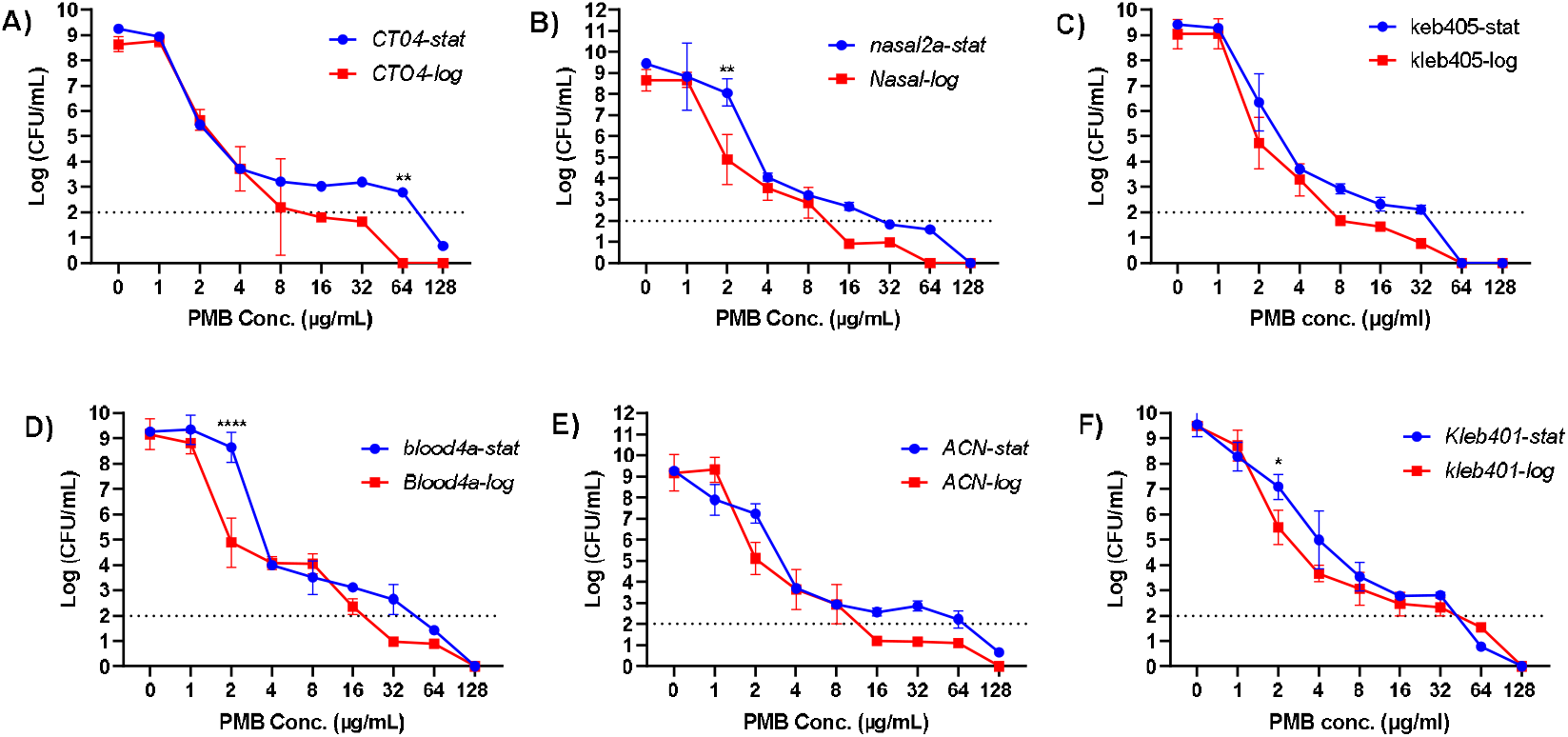
Comparison of heteroresistant subpopulations under log and stationary phase. Three independent PAP experiments were performed from log and stationary phase cultures. Graphs show the log of the mean±SEM CFUs of isolates in the presence of 2-fold increasing concentrations of polymyxin B. The difference between heteroresistance subpopulations under these conditions was statistically determined using two-way ANOVA (Šídák’s multiple comparisons test), with the limit of detection at 100 CFU/ml (log 2 on the y-axis). Significant differences between log and stationary phase cultures for a given concentration are indicated (*).

For each resistant isolate, there was no significant difference between the CFU counts observed at different PMB concentrations relative to no antibiotic control, except at 128 µg/mL where there was a reduction, as determined by the Dunnett’s multiple comparisons test (two-way ANOVA) (**figure 2**). However, whilst there was stable polymyxin resistance, 4/6 isolates (*ACN*^*R*^, Blood4a^R^, *Nasal2a*^*R*^, *CT04*^*R*^) showed morphological instability, with the presence of small colonies on plates with higher PMB concentrations (64 and 128 µg/mL). However, there was no significant difference in the growth profiles of susceptible and heteroresistant populations in the absence of the antibiotic (**Supplementary Figure S1**).

### Characterization of polymyxin B heteroresistance at log and stationary phase

To understand whether the size of the resistant sub-population was influenced by stress, we first examined the impact of growth phase by undertaking PAP experiments for log and stationary phase bacteria. This revealed very similar sized resistant sub-populations in each growth phase. However, there were a few parameters where significant differences were identified for strains *Blood4a* and *CT04*.

### Growth of K. oxytoca in human serum does not influence polymyxin heteroresistance

To examine how the host environment may affect heteroresistance, we first compared the growth of strains and their associated resistant isolates in MHB containing various concentrations of human serum. For all six isolates, both susceptible and heteroresistant populations survived the different serum concentrations ranging from 0.78% to 50%. However, a decrease in bacterial growth was observed with an increase in serum concentration for both populations **(supplementary fig. 2)**. Notably, some of the polymyxin resistant isolates demonstrated reduced growth compared to the susceptible isolates at higher serum concentrations as observed in ACN at 50%, 12.5% and 6.25% serum concentrations (**supplementary fig. 2E)**. Next, we examined the ability of isolates to survive in 100% human serum over 2 hours. All the isolates except *Kleb401* survived at high levels in serum (**Figure 4a**). *Kleb401* was susceptible to serum with complete killing after 90 min in 100% serum.

**Fig. 4:**
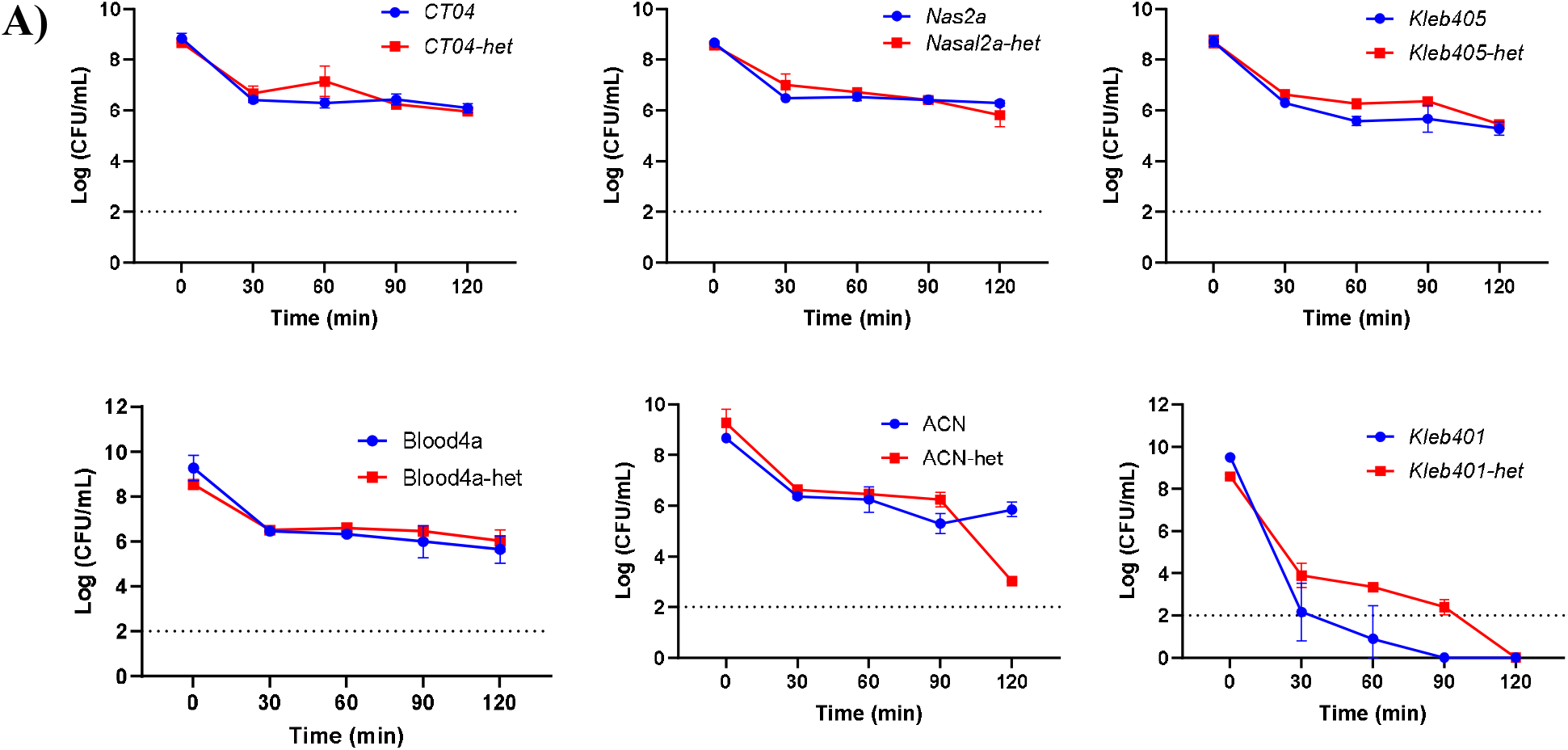
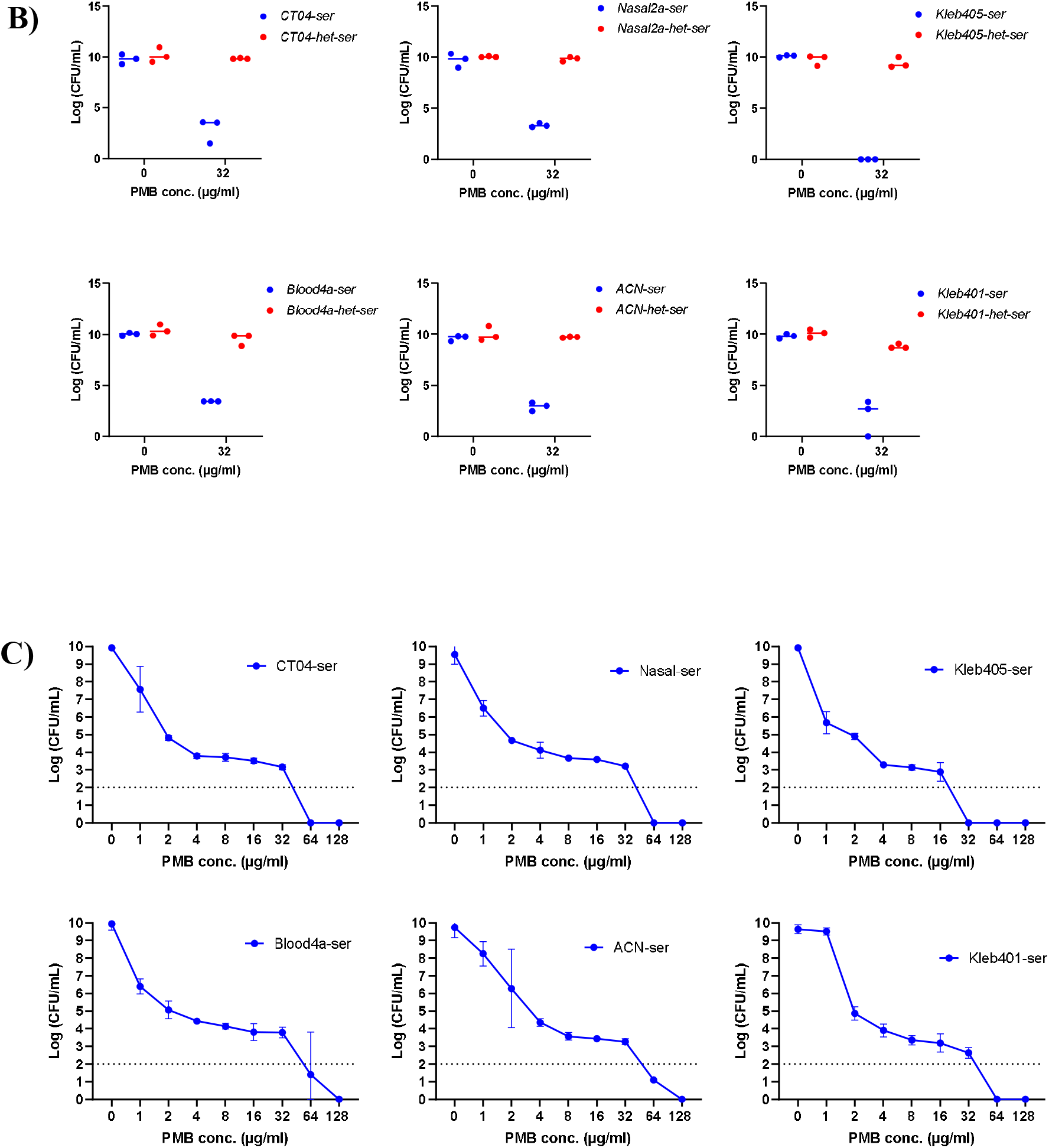
Serum resistance profile of K. oxytoca; **A)** the graph represents three independent time-kill assays of susceptible (*CT04, Nasal2a, Kleb405, Blood4a ACN, Kleb401)* and heteroresistant isolates (*CT04*^*R*^, *Nasal2a*^*R*^ *Kleb405*^*R*^, Blood4a^R^ *ACN*^*R*^, and *Kleb401*^*R*^) grown in 100% human serum for 120 min. Each time-point represents the mean±SEM of CFU count from the three replicates. **B) Polymyxin B susceptibility after growing isolates in serum-adapted MHB**. Susceptible and heteroresistant isolates were grown in 50% serum overnight and plated on PMB (32 µg/mL). The data represent the CFU counts from three independent experiments. **C) Serum has no effect on polymyxin heteroresistance in** *K. oxytoca; the lo*g of the mean±SEM CFUs of three independent biological replicates are shown. The difference between heteroresistance subpopulations from serum-adapted and MHB-grown isolates was statistically determined using two-way ANOVA (Šídák’s multiple comparisons test).

Finally, we examined the effect of human serum on polymyxin B heteroresistance by growing isolates in 50% serum followed by plating on polymyxin B agar plates (32 µg/mL). The heteroresistant population of all six isolates (*Kleb405-het, ACN-het, Blood4a-het, Nasal2a-het, CT04-het*, and *Kleb401-het*) maintained their resistance phenotype **(Figure 4b)**. Similarly, the susceptible population maintained their phenotype, with few resistant colonies observed on PMB (32 µg/mL) compared to the control. We further investigated the effect of serum on polymyxin heteroresistance by subjecting isolates grown in serum to PAP assays **(figure 4c)**. Serum had no significant effect on heteroresistance as determined using two-way ANOVA (Šídák’s multiple comparisons test).

### Lipid A modifications in Polymyxin B resistant subpopulations of K. oxytoca

To understand the underlying mechanism of polymyxin resistance, the Lipid A species present in susceptible and resistant populations of the isolates were profiled by mass spectrometry. Four lipid A species with mass-to-charge ratio (m/z) 1828, 1844, 1909 and 1924 were detected in the susceptible (wild-type) isolates. By contrast, resistant isolates showed two additional peaks centred at m/z 1976 and m/z 1961 which indicates modification of two lipid A species (1844 and 1828) with the addition of L-Ara4N (m/z +131). The proposed structure and component of the native and modified Lipid A species are shown in **figure 5**. We further determined the relative abundance of each Lipid A species in susceptible and resistant subpopulations **(Table 1)**. The levels of native lipid A molecule in susceptible and heteroresistant subpopulations varied with each *K. oxytoca* isolate. For instance, in *CT04*, the levels of lipid A molecule at m/z 1828, 1844, 1909 and 1925 were higher in susceptible populations compared to heteroresistant subpopulations. However, in *Blood4a, Nasal2a* and *Kleb405*, the levels of these four lipid A species were relatively higher in heteroresistant subpopulations. All heteroresistant isolates had significantly high levels of Lipid A species with L-Ara4N modifications **(Table 1)**.

**Table 1:** Relative percentage abundance of native (unmodified) and modified lipid A species in susceptible and resistant subpopulation of *K. oxytoca* isolates.

| Strain |  | LPS modification | (%) Unmodified Lipid A |  |  |  | (%) Modified Lipid A |  |
| --- | --- | --- | --- | --- | --- | --- | --- | --- |
|  |  |  | m/z 1828 | m/z 1844 | m/z 1909 | m/z 1925 | m/z 1961 | m/z 1976 |
| *Susceptible* | *Kleb405* | None | 46.8 | 2.3 | 48.7 | 2.2 | N/A | N/A |
|  | *ACN* | None | 26.3 | 25.7 | 30.0 | 18.1 | N/A | N/A |
|  | *CTO4* | None | 24.4 | 26.9 | 25.3 | 23.3 | N/A | N/A |
|  | *Kleb401* | None | 28.9 | 22.2 | 29.1 | 19.8 | N/A | N/A |
|  | *Blood4a* | None | 24.3 | 19.8 | 32.5 | 23.5 | N/A | N/A |
|  | *Nasal2a* | None | 22.4 | 20.2 | 31.2 | 26.2 | N/A | N/A |
| *Resistant* | *Kleb405<sup>R</sup>* | L-Ara4n | 24.1 | 21.0 | 17.0 | 10.1 | 18.8 | 9.0 |
|  | *ACN<sup>R</sup>* | L-Ara4n | 14.6 | 16.5 | 3.0 | 3.0 | 29.6 | 33.3 |
|  | *CTO4<sup>R</sup>* | L-Ara4n | 14.1 | 16.5 | 14.5 | 16.7 | 14.6 | 23.6 |
|  | *Kleb401<sup>R</sup>* | L-Ara4n | 12 | 35.8 | 10.9 | 16.1 | 9.2 | 16.1 |
|  | *Blood4a<sup>R</sup>* | L-Ara4n | 14.9 | 19.8 | 11 | 16.9 | 13.6 | 23.9 |
|  | *Nasal2a<sup>R</sup>* | L-Ara4n | 12.8 | 16.7 | 25.00 | 13.7 | 11.1 | 20.7 |

N/A \_ Absent

**Fig. 5:**
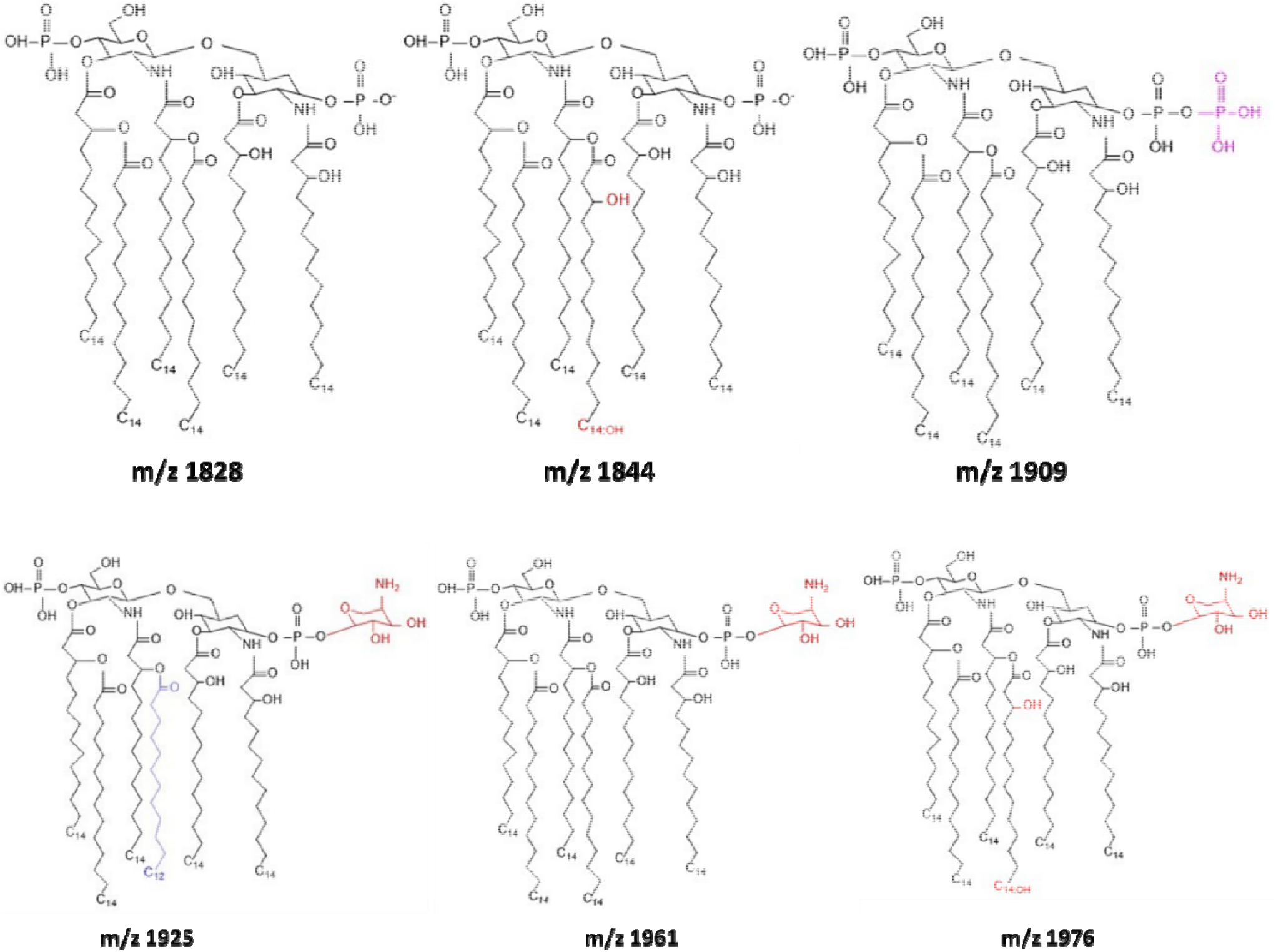
Lipid A profiling of susceptible and heteroresistant isolates showing proposed structures of native and modified lipid A species in susceptible and resistant populations of *K. oxytoca*.

## Discussion

The emergence of multidrug-resistant and heteroresistant *K. oxytoca* isolates in clinical settings is becoming a major public health threat. Importantly, these heteroresistant isolates have frequently been associated with recurrent infections posing a major challenge to antibiotic therapy (Band & Weiss, 2019). In this study, *K. oxytoca* isolates from patient samples and hospital environments were resistant to multiple antibiotics of different classes, including levofloxacin, chloramphenicol, imipenem, and trimethoprim with high MIC values. In previous studies, phylogenetic analysis of diverse *K. oxytoca* isolates has indicated the presence of AMR genes that confer resistance to the classes of antibiotics profiled in this study (Ikhimiukor et al., 2023).

Furthermore, all six isolates profiled in this study demonstrated heteroresistance to polymyxin B, a last resort antibiotic for treating infections caused by MDR Gram-negative bacteria. Heteroresistance was evident through the observation of skipped wells in broth microdilution assays, an indication of heterogeneous bacterial response to antibiotics (Lin et al., 2019). The phenotype was confirmed with PAP assays, the gold standard for detecting antibiotic heteroresistance. Polymyxin heteroresistance has often been reported in Enterobacteriales but with great variability in frequencies across different regions. A three-year study of blood stream isolates of the Enterobacter cloacae complex (ECC) from healthcare centres in Germany reported the widespread of colistin heteroresistance with a frequency of 48.4% and MIC range between 64-512 µg/mL (Doijad et al., 2023). In a large U.S. surveillance of 408 carbapenem-resistant Enterobacterales isolates collected between 2012–2015, colistin heteroresistance was detected in 10.1% of isolates although 93.2% of those were misclassified as susceptible by routine tests (Band et al., 2021). This highlights the tendencies of the phenotype to escape clinical detection. The detection of heteroresistance in all our isolates therefore supports evidence of the emergence of heteroresistance in clinical isolates and confirms the need for a more robust detection method especially in healthcare settings.

Currently, reports on polymyxin heteroresistance in *Klebsiella* species primarily focus on *K. pneumoniae*. The first U.S. report of polymyxin heteroresistance in *Klebsiella* species was demonstrated in two distinct isolates of *K. pneumoniae*, where resistant subpopulations had colistin MICs up to 100 µg/mL while the main population remained susceptible at 2 µg/mL (Band et al., 2018). Similarly, Halaby et al. (2016) reported colistin heteroresistance in ESBL-producing *K. pneumoniae* clinical isolates, with resistant subpopulation MICs ranging from 8 to 64 µg/mL as determined by population analysis profiles (PAP)). In this study, however, we focused on characterization of polymyxin heteroresistance in *K. oxytoca*, which is often under-recognized due to the lack of routine speciating methods in clinical diagnostics. We observed similar heteroresistance phenotypes, with high MICs of resistant subpopulation ranging from 32 to 128 µg/mL. The emergence of heteroresistance in these *K. oxytoca* isolates suggests their ability to cause persistent or recurrent infections as indicated in *in vivo* experiment with heteroresistant *K. pneumoniae* isolates (Band et al., 2018). The isolates profiled in this study showed a stable heteroresistance phenotype such that, a single colony of each isolate produces a small highly resistant subpopulation. This phenotype was not affected by bacterial or host environmental factors such as log phase, stationary phase and human serum.

Polymyxin B is a membrane-targeting cationic antibiotic that primarily interacts with the negatively charged LPS of the bacterial membrane. Polymyxin resistance has been associated with structural modification of the LPS such as the addition 4-amino-L-arabinose (L-Ara4N) or phosphoethanolamine (pEtN) to the lipid A component (Dortet et al., 2020). These modifications have also been linked to colistin resistance and heteroresistance in *K. pneumoniae* and *E. cloacae* respectively (Kang et al., 2019). Hence, we profiled the lipid A of all six *K. oxytoca* isolates to identify modifications that may be implicated in the observed phenotype. Four native lipid A species were identified in the susceptible population of the six isolates. In the resistant subpopulations, however, these native lipid A species were modified by the addition of an L-Ara4N moiety, consistent with previous observations in *K. pneumoniae*

This study is the first to characterize polymyxin heteroresistance in *K. oxytoca* as well as factors that could potentially influence the phenotype. These isolates exhibit a stable heteroresistance which is not influenced by bacterial or host environmental factors. This phenotype suggests an inherent ability to survive antibiotics and cause persistent infections. Additionally, resistant subpopulations had L-Ara4N-modified lipid A, a potential mechanism driving polymyxin heteroresistance in *K. oxytoca*. As an emerging threat in healthcare settings, we suggest further studies to explore the mechanisms driving heteroresistance to polymyxins and other antibiotics to inform appropriate therapeutic strategies.

## Acknowledgements

The authors would like to acknowledge the members and staff of the AMR Research Group (led by AI) at the West African Centre for Cell Biology of Infectious Pathogens (WACCBIP) University of Ghana for technical support.

