## Supplementary data file for "Polymyxin heteroresistance in *Klebsiella oxytoca*"

**Table S1:** Antimicrobial resistance patterns of *K. oxytoca* isolates

| **Antibiotics** | **Minimum inhibitory concentration (µg/mL)**  **Bacterial strains** | | | | | |
| --- | --- | --- | --- | --- | --- | --- |
|  | ***Kleb401*** | ***Nasal2A*** | ***CT04*** | ***Blood4a*** | ***can*** | ***Kleb405*** |
| **Ceftazidime** | 0.25 | 2 | 2 | 2 | 1 | 0.5 |
| **Levofloxacin** | 64 | 64 | 64 | 64 | 32 | 32 **(s.w)** |
| **Chloramphenicol** | >64 | >64 | >64 | >64 | >64 | 64 |
| **Colistin** | 16 **(s.w)** | 2 | 32 **(s.w)** | 64 **(s.w)** | 32 **(s.w)** | 8 **(s.w)** |
| **Polymyxin B** | 16 **(s.w)** | 32 **(s.w)** | 16 **(s.w)** | 32 | 16 **(s.w)** | 0.25 |
| **Gentamicin** | 0.25 | 0.25 | 0.25 | 0.25 | 0.2 | 0.25 |
| **Imipenem** | 16 | 16 | 32 | 32 | 32 | 16 |
| **Meropenem** | 0.125 | 0.125 | 0.125 | 0.125 | 0.125 | 0.125 |
| **Trimethoprim** | >64 | >64 | >32 | >64 | >64 | 32 **(s.w)** |

***s.w_ skipped wells**

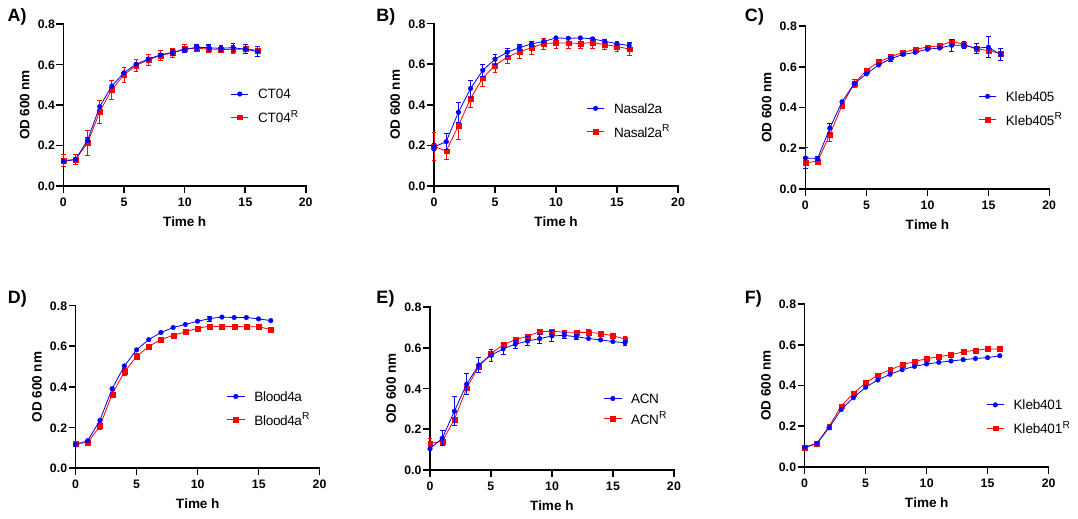

**Supplementary figure S1:** Growth profiles of susceptible and heteroresistant subpopulations of all six isolates. The mean±SEM OD600 values from three independent biological replicates is shown.

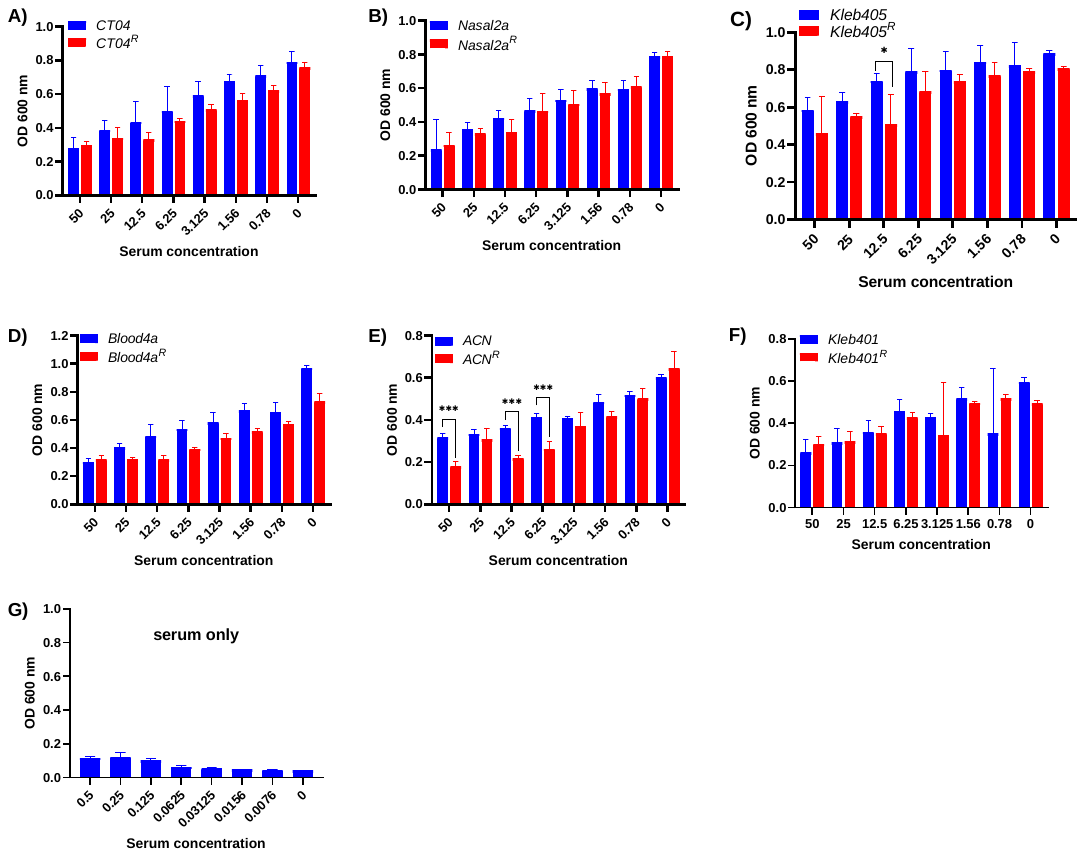

**Supplementary figure S2: Serum growth assay.** Growth of susceptible and resistant isolates in MHB containing various concentrations of serum. The graphs show the mean±SEM of OD measurement from three independent experiments. Statistical difference between bacterial survivals at different serum concentrations compared to the control was determined using two-way ANOVA (Šídák's multiple comparisons test).

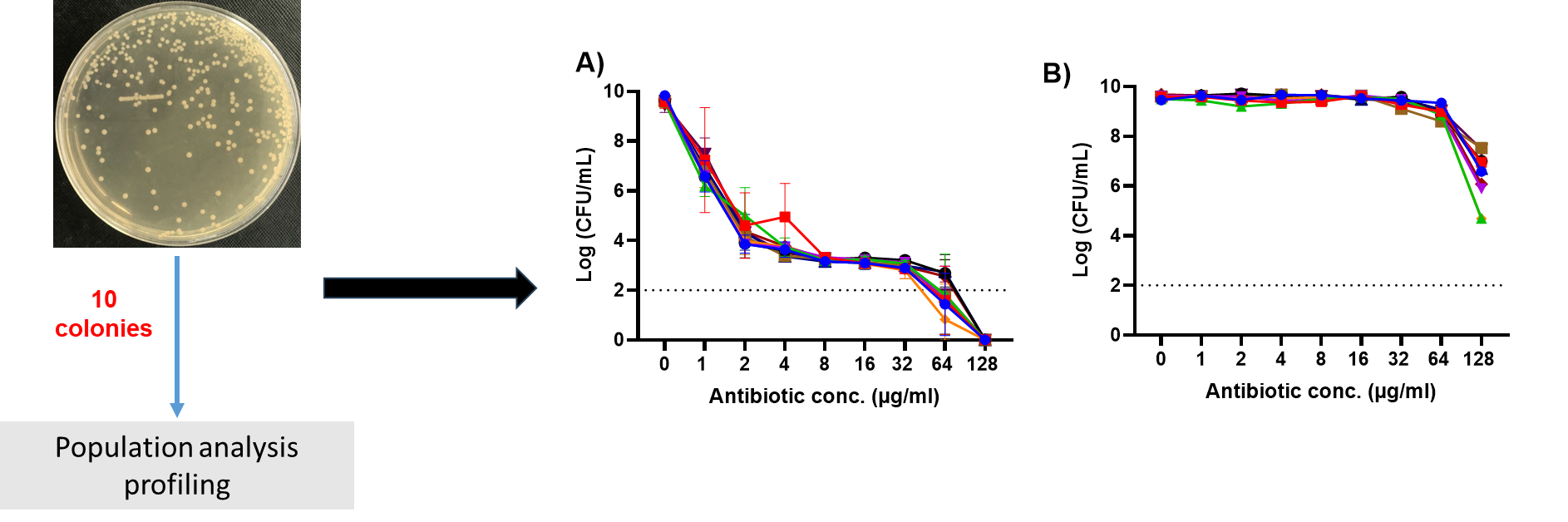

**S.3: Determination of the stability of heteroresistance in *K. oxytoca (ACN*);** ten colonies of the isolate (ACN) were randomly selected from MH agar plate and subjected to PAP experiment **(A)**. Heteroresistant isolates obtained from MH agar containing 32 µg/Ml of polymyxin B were taken through another round of PAP experiment **(B)**. The graphs show the mean±SEM of three independent experiments.

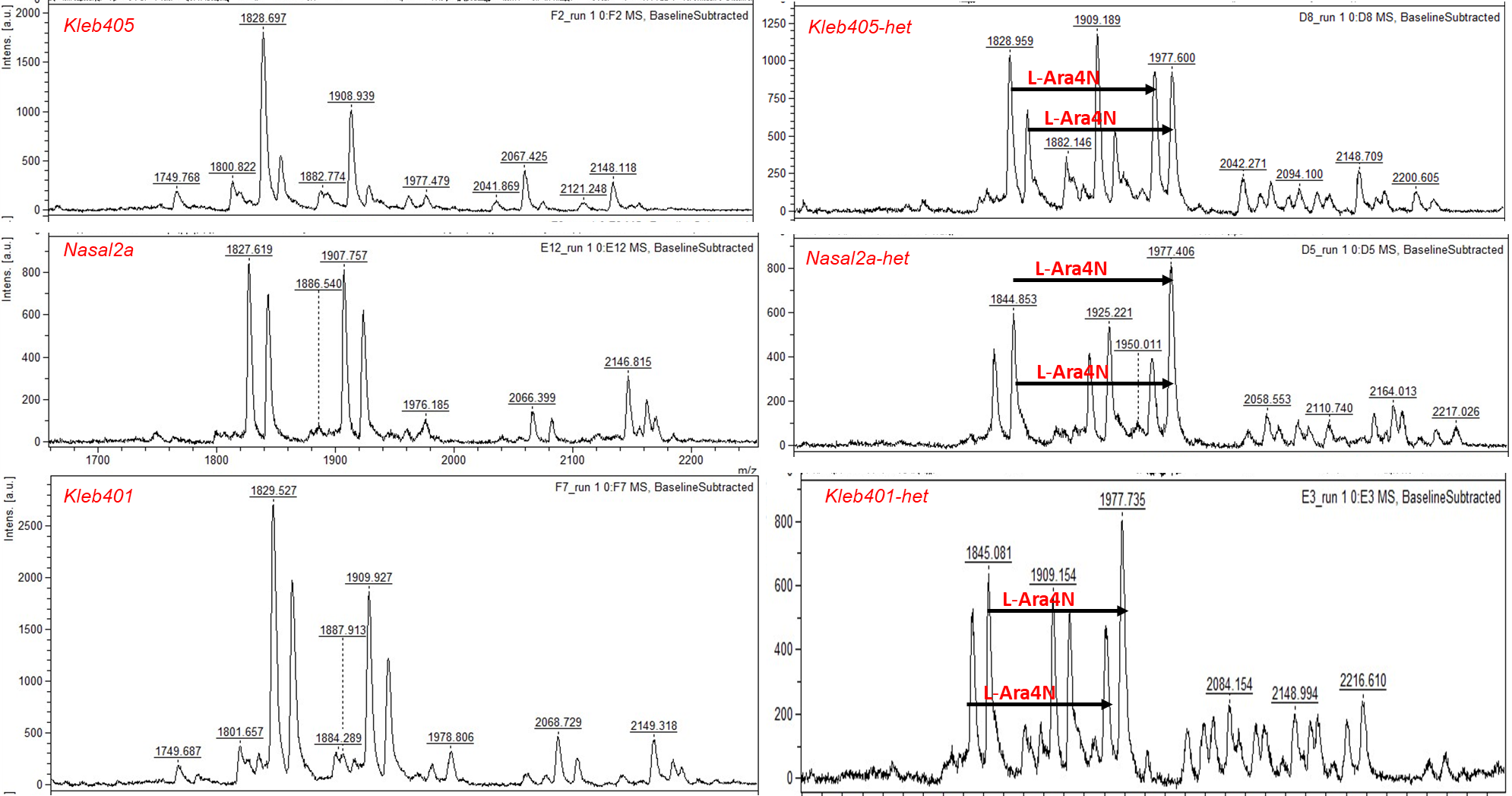

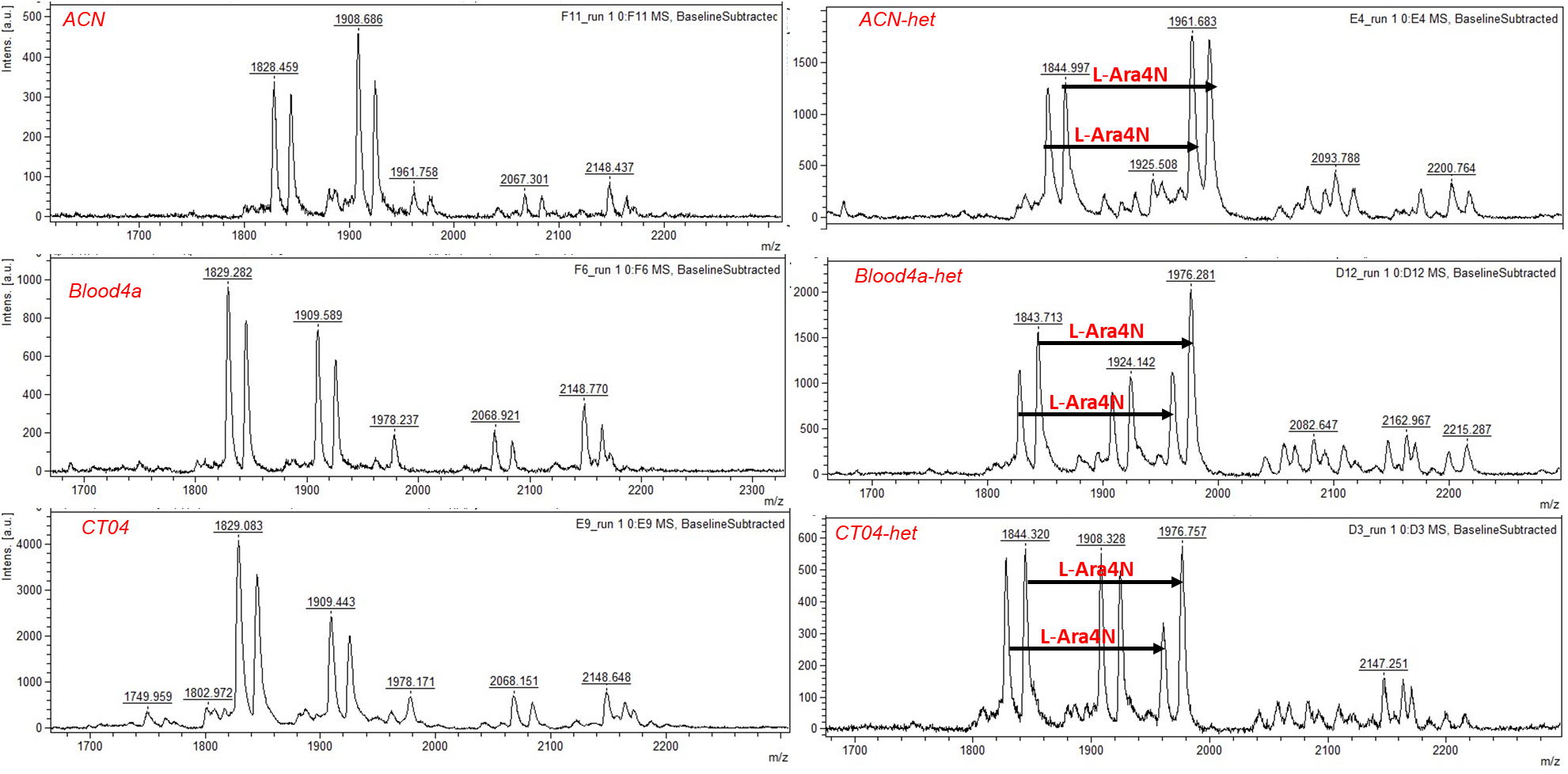

**S.4:** Representative mass spectra of polymyxin susceptible and heteroresistant subpopulations of all six isolates *(Kleb405, Nasal2a, Kleb401, ACN, Blood4a, and CT04)*
